# Prevalence of loss-of-function, gain-of-function and dominant-negative mechanisms across genetic disease phenotypes

**DOI:** 10.1101/2025.03.13.642984

**Authors:** Mihaly Badonyi, Joseph A Marsh

## Abstract

Molecular disease mechanisms caused by mutations in protein-coding regions are diverse, but they can be broadly categorised into loss-of-function (LOF), gain-of-function (GOF), and dominant-negative (DN) effects. Accurately predicting these mechanisms is a pressing clinical need, as therapeutic strategies must align with the underlying disease mechanism. Moreover, computational predictors tend to perform less well at the identification of pathogenic GOF and DN variants. Here, we develop a protein structure-based missense LOF (mLOF) likelihood score that can separate recessive LOF and dominant LOF from alternative disease mechanisms. Using mLOF scores, we estimated the prevalence of molecular mechanisms across 2,837 phenotypes in 1,979 Mendelian disease genes, finding that DN and GOF mechanisms account for 48% of phenotypes in dominant genes. Applying mLOF scores to genes with multiple phenotypes revealed widespread intragenic mechanistic heterogeneity, with 43% of dominant and 49% of mixed-inheritance genes harbouring both LOF and non-LOF mechanisms. Furthermore, we show that combining mLOF scores with phenotype semantic similarity enables the prioritisation of DN mechanisms in mixed-inheritance genes. Our structure-based approach, accessible via a Google Colab notebook, offers a scalable tool for predicting disease mechanisms and advancing personalised medicine.

## Introduction

The vast majority of disease-causing genetic variants identified to date are located within protein-coding regions of the genome. While many lead to a loss of protein function (LOF), often through premature stop codons or missense changes that destabilise protein folding, others exert their effects via alternative (non-LOF) mechanisms, such as gain-of-function (GOF) or dominant-negative (DN) effects. Understanding these mechanisms usually requires examining how mutant proteins interact with other molecules. Their impacts can manifest through various means, including disruption or creation of novel interactions, altered binding affinity or specificity, changes to protein complex assembly, and induction of aggregation, mislocalisation, or phase separation^1^. The diversity of molecular mechanisms presents a significant challenge for their identification, often necessitating elaborate experimental strategies to validate them^2^, which are costly and time-consuming.

Accurate prediction and validation of molecular disease mechanisms are essential for developing effective targeted therapies. Diseases resulting from LOF mutations are usually amenable to gene therapy, where the delivery of functional gene copies compensates for the defective allele. This approach has successfully treated conditions such as RPE65-associated retinal dystrophy^3^ and Duchenne muscular dystrophy^4^. In contrast, diseases caused by non-LOF mutations are more suited to treatment with small molecules that inhibit the altered or excessive function, as demonstrated by the development of KRAS degraders for cancer^5^, or through gene-editing and silencing strategies, as exemplified by a promising treatment for retinitis pigmentosa, driven by the GOF mutation p.Pro23His in rhodopsin^6^. Similar allele-specific targeting approaches offer hope for treating DN conditions, such as collagen-related dystrophy^7^ and long QT syndrome^8^. While most genes are associated with a single molecular mechanism, some are known to exhibit multiple mechanisms, requiring distinct therapeutic interventions. For example, sodium channel blockers are effective for epilepsy associated with GOF variants in SCNA1^9^, whereas gene replacement therapy may soon address SCNA1 haploinsufficiency in Dravet syndrome^10^.

Despite the clear clinical need, predicting molecular disease mechanisms remains difficult. Current computational methods usually focus on predicting LOF and function-altering mechanisms at the level of individual genetic variants^11–13^. However, there are also gene-level features that tend to be associated with different mechanisms^14–17^. We recently developed a model to predict the most likely mechanism when heterozygous disease mutations are found in a gene^18^. These predictions have now been incorporated into the DECIPHER database^19^, assisting clinicians in identifying potential disease mechanisms.

We previously reported two structural properties—specifically, the energetic impact and clustering of missense variants—that discriminate between genes with LOF and non-LOF mechanisms exceptionally well^15^. This is because LOF mutations tend to be highly destabilising and spread throughout protein structures, whereas non-LOF mutations, which are structurally milder, often exhibit clustering within functionally important regions. We quantify the impacts of variants on protein stability using changes in Gibbs free energy of folding (ΔΔG) predicted with FoldX^20^, while variant clustering is assessed with the extent of disease clustering (EDC) metric^15^. While ΔΔG is calculated at the variant level, EDC operates at an intermediate level, requiring multiple variants but not necessarily all disease variants within a gene. This flexibility enables EDC to be applied to a group of variants, particularly those associated with the same phenotype, as demonstrated for cancer-associated and Weaver syndrome variants in EZH2^21^.

In this study, by integrating EDC and ΔΔG data from pathogenic variants, we develop an empirical distribution-based method to derive a missense LOF (mLOF) likelihood score, and demonstrate its utility for improving molecular mechanism predictions in a Bayesian framework. By assembling phenotype annotations for over 70% of pathogenic missense variants in ClinVar^22^, we show that the mLOF score is particularly powerful at the phenotype level. Most importantly, we estimate the prevalence of molecular mechanisms across genetic disease phenotypes, revealing widespread mechanistic heterogeneity and highlighting its implications for precision medicine. We make our method available as a Google Colab notebook, allowing mLOF score calculation for variant sets in human protein-coding genes at https://github.com/badonyi/mechanism-prediction.

## Results

### Developing the mLOF score for predicting missense variant molecular mechanisms

Our objective was to predict the likelihood of a set of missense variants being associated with LOF *vs* non-LOF molecular mechanisms by integrating information about their protein structural context. Specifically, we sought to combine clustering in three-dimensional space, as quantified by EDC, and predicted energetic impacts, as measured by ΔΔG. To achieve this, we developed an approach based on the empirical distributions of these metrics in LOF and non-LOF genes^15^, *i.e*., genes with pathogenic missense variants known to act via LOF and DN or GOF mechanisms, respectively. Importantly, we use ΔΔG_rank_ in place of raw ΔΔG values. This is a recently introduced rank-normalised metric that improves interpretability and facilitates comparisons across different proteins^23,24^. For a given observation of EDC and ΔΔG_rank_ in a set of variants, we calculate the marginal probabilities of these observations being drawn from the LOF rather than non-LOF distributions (**Figure S1**). The probabilities are then combined into the mLOF score, which represents the likelihood that the variants will have a LOF effect given their energetic impact and dispersal within the protein structure.

To evaluate the utility of the mLOF score, we treated predictions from our previously published proteome-scale model (pDN/GOF/LOF) as informative priors for the likelihood of a disease mechanism occurring in a gene^18^. By updating these priors with the mLOF score, we derived mechanism-specific posterior scores (postDN/GOF/LOF), which represent adjusted estimates of the likelihood that a gene exhibits a mechanism, taking into account the structural properties of its pathogenic missense variants. **Figure 1A** provides a graphical overview of our method.

**Figure 1.**
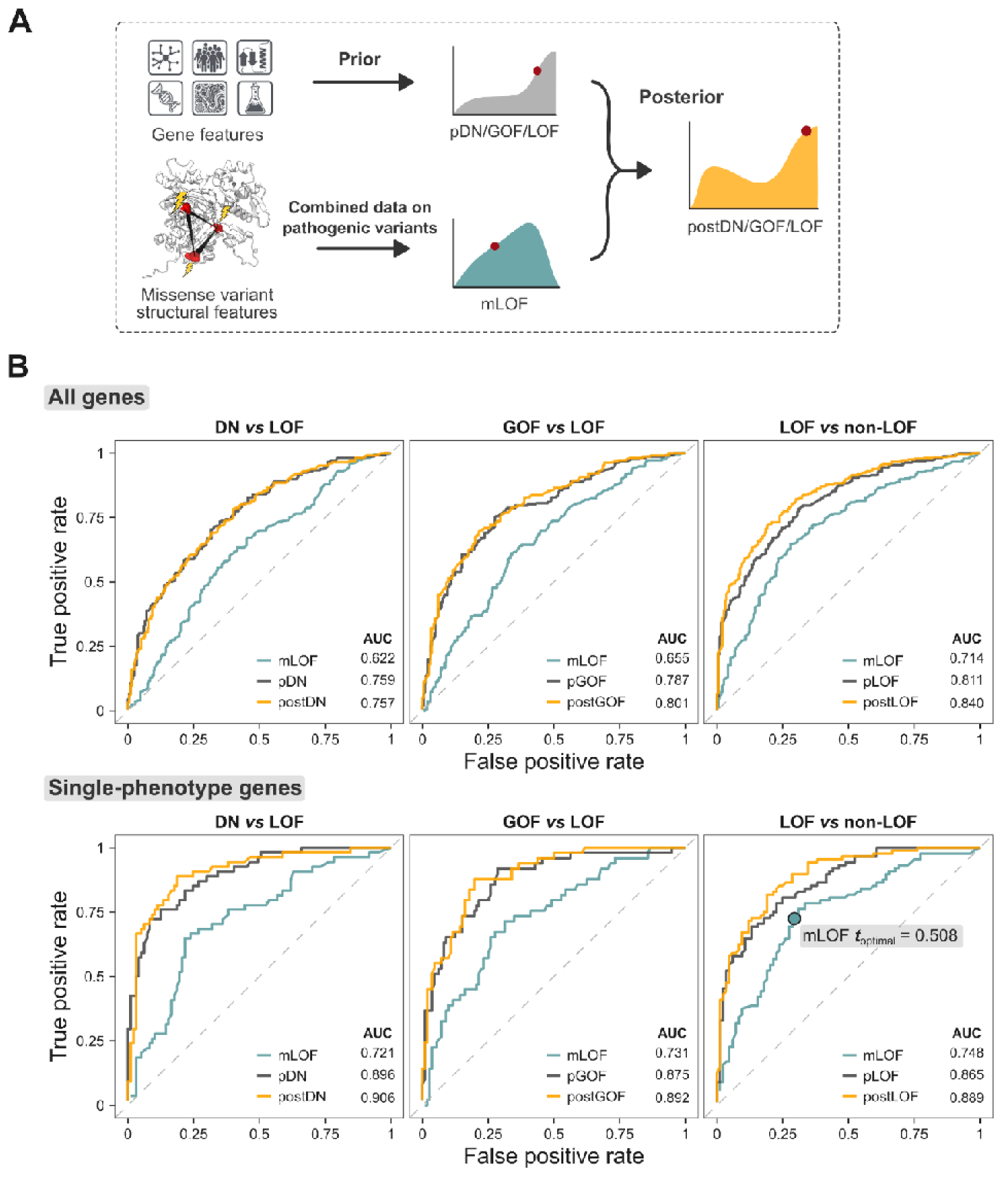
Predicting the likelihood of a loss-of-function mechanism based on the structural properties of pathogenic missense variants. **A:** Overview of the mLOF score framework. The missense LOF likelihood score mLOF is calculated from empirical distributions of the metrics EDC (spatial clustering) and ΔΔG_rank_ (energetic impact) in LOF and non-LOF genes. This score is then used to update gene-level mechanism-specific priors (pDN/GOF/LOF) established in an earlier study^18^. The final posterior scores (postDN/GOF/LOF) represent adjusted estimates of the likelihood that a gene exhibits a specific molecular disease mechanism, given the structural properties of its pathogenic missense variants. **B:** Receiver operating characteristic (ROC) curves and area under the curve (AUC) values of the mLOF score, the prior mechanism probability for the gene, and the posterior mechanism-specific scores across the binary class pairs used to construct the priors. The analysis is split into all genes, using all pathogenic missense variants, and a subset of single-phenotype genes, where only variants linked to the specific OMIM phenotypes are considered. mLOF ***t***_optimal_ shows the optimal ROC threshold.

We first applied this method to pathogenic missense variants in exclusively autosomal dominant (AD) genes with gene-level molecular mechanism classifications^18^, and calculated the area under the receiver operating characteristic curve (AUROC) for the mLOF score, as well as the prior and posterior mechanism-specific scores (**Figure 1B**). We found the mLOF score to be predictive across the binary class pairs previously used to construct the priors (DN *vs* LOF, GOF *vs* LOF, and LOF *vs* non-LOF), with AUROC ranging from 0.622 to 0.714, indicating generalisability across the mechanisms.

One possible explanation for the limited performance is that many genes are associated with multiple molecular disease mechanisms, which imposes fundamental limitations on our gene-level approach. Although we only have gene-level rather than phenotype-level classifications, one way of addressing this limitation is by considering those genes with a single disease phenotype, which are thus more likely to be associated with a unique mechanism. Therefore, we used variant-level phenotype annotations from the Online Mendelian Inheritance in Man (OMIM) database^25^ to identify dominant genes associated with a single disease phenotype.

Notably, AUROC values were markedly increased across all binary class pairs (**Figure 1C**). A similar conclusion is supported by the area under the balanced precision-recall curve analysis^26^ (**Figure S2**). We also derived the optimal threshold for distinguishing between LOF and non-LOF mechanisms using the single phenotype genes. The resulting value of 0.508 provides a practical cutoff for assessing whether a group of variants is likely to exhibit a LOF mechanism and can be used to compare different variant groups in the same gene.

As an additional validation, we applied the mLOF score to previously published high-throughput functional assay data from TP53^27^ and HRAS^28^, evaluating its capacity to distinguish LOF, DN, and GOF variants within the same gene. For TP53, we found that variants with a LOF mechanism had the highest mLOF score (0.551), primarily driven by the dispersal of variants in the structure (**Figure S3A**). DN variants, in contrast, had a lower mLOF score of 0.445 and were concentrated within the DNA-binding domain. Notably, variants exhibiting both DN and LOF properties in the assay clustered exclusively in the DNA-binding domain, showed the highest predicted structural destabilisation, and had the lowest mLOF score (0.351). We speculate that these variants are highly destabilising in TP53 knockout assays, but may achieve partial stabilisation through wild-type binding, thus manifesting a DN effect in a context-dependent fashion. In the HRAS dataset, a clear difference was observed between GOF and LOF variants, with mLOF scores of 0.426 and 0.613, respectively (**Figure S3B**). GOF variants were clustered at key functional sites, whereas LOF variants spread across protein core residues.

Furthermore, we evaluated the mLOF score against GOF predictions by the LoGoFunc method^11^. Although LoGoFunc provides GOF probabilities at the variant level, averaging these probabilities for a phenotype yields a measure comparable to the mLOF and postGOF scores. We tested the performance of this metric in dominant single-phenotype genes, using both all available genes and the test set of our gene-level predictor. As shown in **Figure S3C-D**, in both cases, when combined with the prior GOF mechanism likelihood for the gene, mLOF yielded postGOF scores that substantially outperformed the average GOF probabilities from LoGoFunc.

### Prevalence of molecular mechanisms across disease phenotypes

Motivated by these findings, we set out to assess the prevalence of molecular mechanisms across genetic disease phenotypes. We first classified disease phenotypes on the basis of their inheritance. Specifically, genes can show either exclusively autosomal dominant (AD) or autosomal recessive (AR) inheritance, or they may show mixed inheritance, being associated with both dominant and recessive variants. Dominant and recessive variants in mixed-inheritance genes may be associated with distinct phenotypes, in which case we can consider the dominant ([AD]-AD/AR_mixed_) or recessive ([AR]-AD/AR_mixed_) phenotypes separately. In contrast, as we only have gene-level phenotype:inheritance associations available from OMIM, for those genes with mixed-inheritance associated with the same phenotype, we are unable to distinguish between dominant and recessive variants so they are considered together (AD/AR_same_). We also classified AD phenotypes on the basis of molecular mechanisms, using our previous gene-level LOF, GOF and DN annotations. These phenotype classifications are summarised in **Table 1**.

**Table 1.** Inheritance- and mechanism-based classification of disease phenotypes.

| **Inheritance-based phenotype classification** | **Abbreviation** |
| --- | --- |
| Phenotypes of exclusively autosomal recessive genes | AR |
| Autosomal recessive phenotypes of mixed-inheritance genes | [AR]-AD/AR<sub>mixed</sub> |
| Autosomal dominant phenotypes of mixed-inheritance genes | [AD]-AD/AR<sub>mixed</sub> |
| Phenotypes of autosomal genes inherited in both dominant and recessive modes | AD/AR<sub>same</sub> |
| Recessive phenotypes of X-linked genes | XLR |
| Phenotypes of exclusively autosomal dominant genes | AD |
| **Mechanism-based phenotype classification** |  |
| Autosomal dominant phenotypes in genes with a loss-of-function mechanism | LOF |
| Autosomal dominant phenotypes in genes with a gain-of-function mechanism | GOF |
| Autosomal dominant phenotypes in genes with a dominant-negative mechanism | DN |
| Autosomal dominant phenotypes in genes without a reported molecular mechanism | Unknown |

**Figure 2A** shows the mLOF score distribution for the different inheritance-based phenotype classifications, ordered by their mean. The observed distributions align remarkably well with our expectations: AR, [AR]-AD/AR_mixed_ and XLR phenotypes display the highest mLOF scores. AD phenotypes, while shifted to the left side of the optimal threshold by the mean, are evidently bimodal, suggesting the presence of both LOF and non-LOF mechanisms. Interestingly, [AD]-AD/AR_mixed_ phenotypes fall on the left side of the optimal threshold, with a mean of 0.499. This can be explained by considering that the coexistence of a recessive disorder provides a level of evidence against dosage sensitivity^29^, thus making AD phenotypes in these genes more likely to arise through alternative mechanisms. In contrast, AD/AR_same_ phenotypes have a higher mean mLOF score relative to AD phenotypes, which could result from the unequal mixing of AD and AR variants—a limitation inherent to our use of phenotype-level rather than variant-level annotations. Alternatively, missense variants in these phenotypes may follow a single inheritance mode, while other mutation types, such as stop-gain or frameshift mutations, correspond to the other mode. This phenomenon has been observed, for example, in *ITPR1*, where homozygous null and *de novo* missense variants both cause Gillespie syndrome^30^.

**Figure 2.**
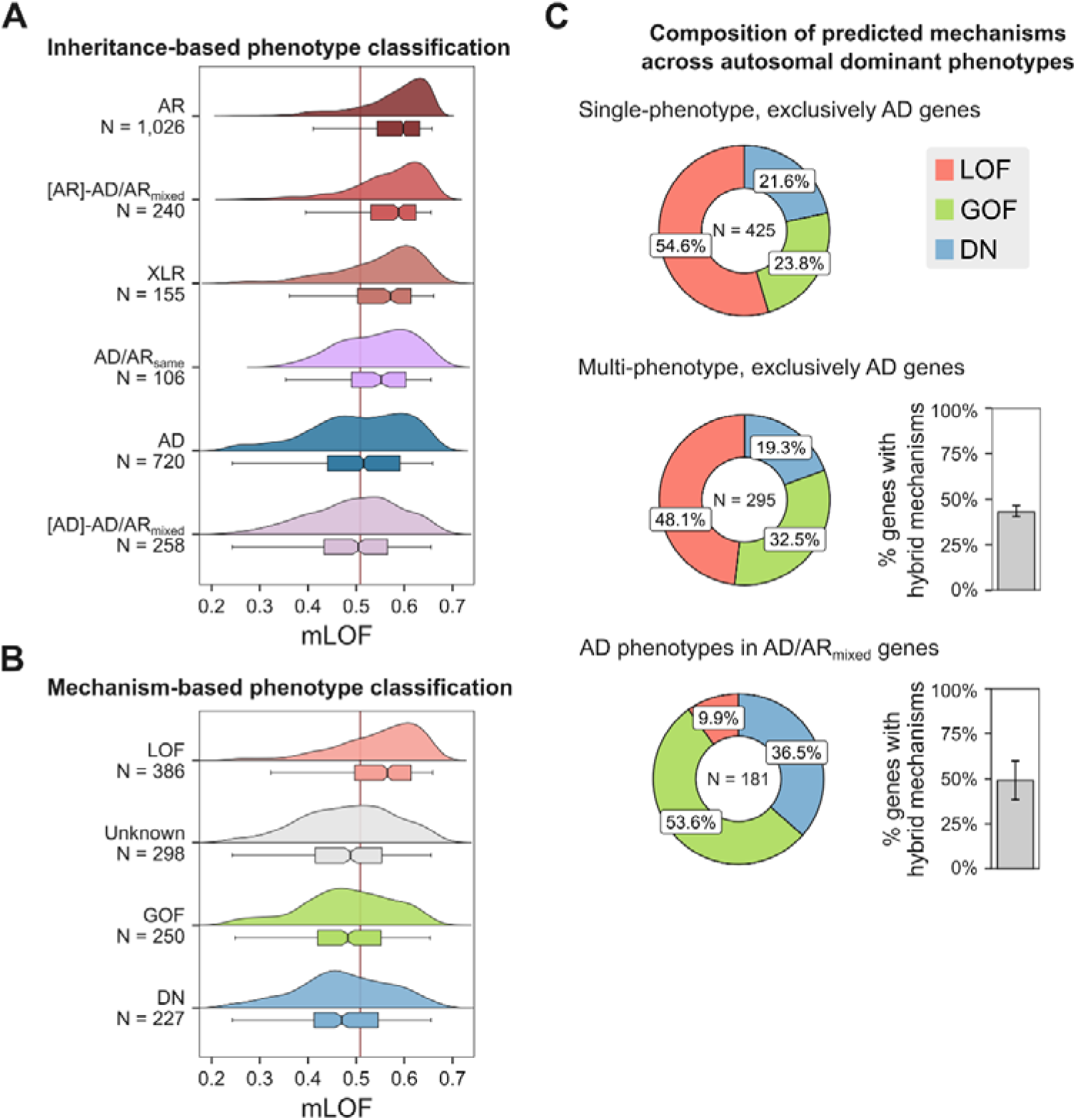
mLOF scores reveal the prevalence of molecular mechanisms at the phenotype level. **A-B**: Distribution of mLOF scores for inheritance- and mechanism-based phenotype classifications (see **Table 1** for the description of the abbreviations). N denotes the number of phenotypes in each group. Red line indicates the optimal mLOF score threshold. **C**: The fractional composition of predicted mechanisms for phenotypes across the indicated categories. The predictions represent the highest-ranking posterior mechanism-specific score for each phenotype. N denotes the number of phenotypes in each group. Bar charts show estimates of genes with both LOF and non-LOF (‘hybrid’) mechanisms. Error bars are 95% credible intervals calculated from a posterior distribution of fractions derived using the bootstrap estimates of the optimal mLOF threshold.

The different mechanism-based phenotype classifications are shown in **Figure 2B**. As expected, dominant LOF phenotypes have the highest mean mLOF score (0.547), while GOF and DN phenotypes are strongly left-shifted, with mean mLOF scores of 0.480 and 0.474, respectively. Unknown phenotypes, those of dominant genes without reported mechanisms, show a left-skewed distribution with a mean of 0.484. This likely reflects detection bias, as non-LOF variants are more difficult to experimentally characterise and less well predicted by computational tools, leading to an apparent enrichment of alternative mechanisms in these genes.

Next, we classified AD phenotypes based on their highest mechanism-specific posterior scores into LOF, GOF, and DN categories to assess the contribution of different molecular mechanisms. We focused on three groups in particular: exclusively AD genes with a single phenotype, those with multiple phenotypes, and AD phenotypes in mixed-inheritance genes, *i.e*., genes associated with both AD and AR disorders. In **Figure 2C**, we show the composition of predicted molecular mechanisms across these groups. Single-phenotype AD genes exhibited the largest fraction of phenotypes with a LOF mechanism, at 54.6%. The remaining fraction was attributed to GOF and DN mechanisms occurring at similar frequencies, at 23.8% and 21.6%, respectively. In multi-phenotype AD genes, the fraction of phenotypes with a LOF mechanism was lower, at 48.1%, followed by GOF at 32.5% and DN at 19.3%. This difference may be explained by the observation that multiple disease phenotypes are unlikely to occur in the same gene due to haploinsufficiency; thus, by exclusion, a greater contribution of non-LOF mechanisms is expected. AD phenotypes in mixed-inheritance genes had the lowest proportion of LOF mechanisms, at just 9.9%, followed by GOF at 53.6% and DN at 36.5%. As observed with the mLOF score distribution of these genes in **Figure 2A**, this likely reflects a reduced likelihood of haploinsufficiency conferred by the presence of a recessive disorder, which makes dominant phenotypes more likely to arise through alternative mechanisms.

We next estimated the fraction of multi-phenotype genes with disease phenotypes involving both LOF and non-LOF molecular mechanisms. This analysis revealed that 43.1% of multi-phenotype AD genes accommodate at least one DN or GOF disease mechanism in addition to LOF. Similarly, in mixed-inheritance genes, we estimated a frequency of 49.1%, assuming most recessive disorders involve biallelic LOF (with rare exceptions^31,32^), and quantifying the fraction with a dominant non-LOF mechanism. These findings suggest that, based on the structural properties of missense variants, mechanistic heterogeneity is widespread among multi-phenotype genes. To facilitate access to these results, we provide a comprehensive list of OMIM phenotypes (*N* = 2,837) in **Table S1**, including MIM identifiers, disease names, EDC and ΔΔG_rank_ values, mLOF scores, and the mechanism-specific posterior scores.

### Dominant-negative phenotypes in mixed-inheritance genes

Intriguingly, our results suggest that LOF is very rare as a mechanism underlying dominant phenotypes in mixed-inheritance genes, accounting for only 9.9% of cases (**Figure 2C**). While this might in part be explained by considering that mixed-inheritance genes are less likely to be haploinsufficient, there are many examples where the same phenotype is associated with both dominant and recessive variants. One possible explanation is that the recessive variants are hypomorphic, causing only a partial LOF in each allele that amount to the same net wild-type activity level as complete LOF in one allele. To test this hypothesis, we compared ΔΔG_rank_ distributions of recessive phenotypes in mixed-inheritance genes ([AR]-AD/AR_mixed_) with those in exclusively AR genes (**Figure 3A**). We observed that [AR]-AD/AR_mixed_ phenotypes exhibit lower ΔΔG_rank_ values compared with those of AR genes (*P* = 1.6 × 10^-3^, Wilcoxon rank-sum test), consistent with the presence of hypomorphic variants. Unfortunately, we do not have variant-level inheritance classifications in AD/AR_same_ genes, where we might expect the tendency for recessive variants to be hypomorphic to be even greater, thus explaining the equivalent phenotypes for dominant and recessive variants. Nonetheless, the case of *PKD1*, where recessive hypomorphic variants have recently been implicated in polycystic kidney disease^33^— the same phenotype for which there is sufficient evidence of haploinsufficiency caused by dominant LOF mutations (ClinGen^34^ Curation ID: 007675)—underscores the relevance of these effects.

**Figure 3.**
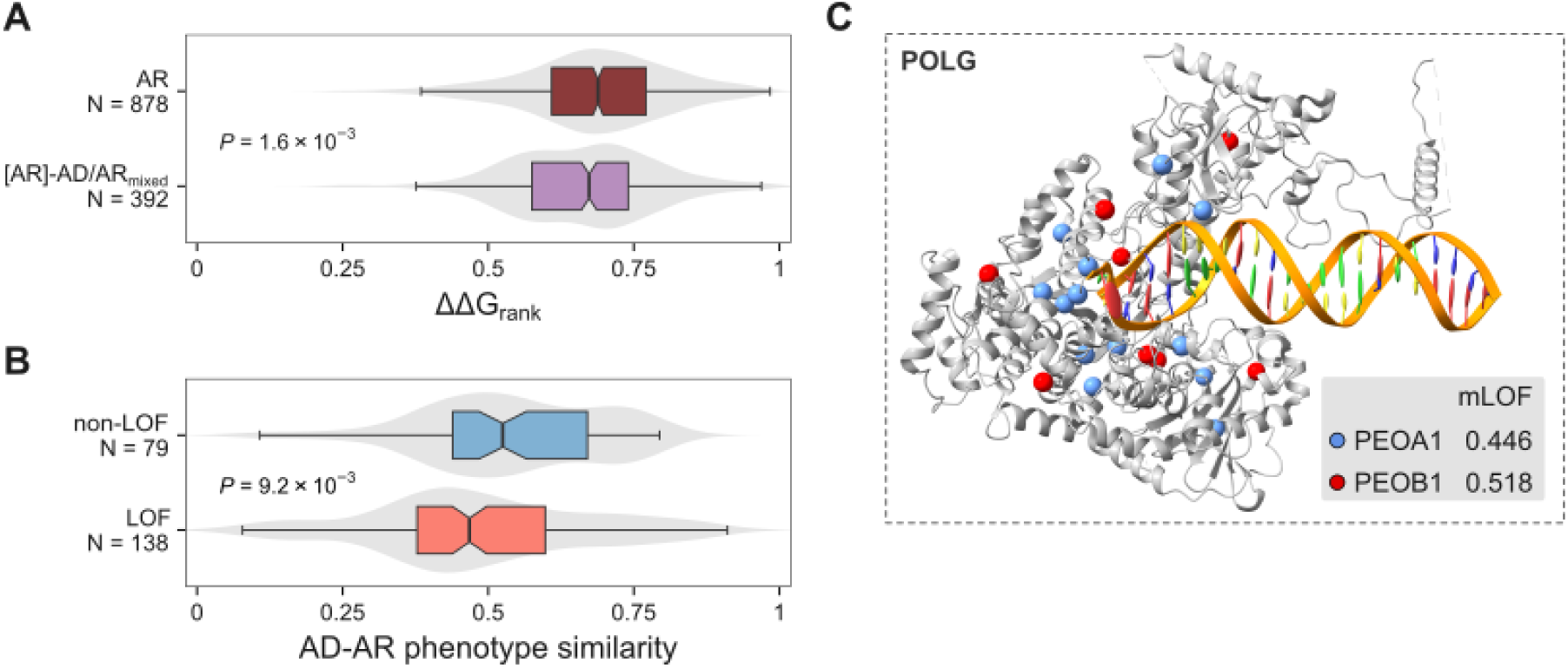
mLOF score and phenotype semantic similarity prioritise dominant-negative phenotypes in AD/AR_mixed_ genes. **A**: ΔΔG_rank_ distributions for disease phenotypes in exclusively recessive genes (AR) and recessive phenotypes in AD/AR_mixed_ genes ([AR]-AD/AR_mixed_). The *P*-value represents a two-sided Wilcoxon rank-sum test. **B**: Comparison of AD-AR phenotype semantic similarity within [AR]-AD/AR_mixed_ genes, split into whether the variants of the AD phenotype are predicted to have a non-LOF effect. **C**: *POLG* (AlphaFold model superimposed onto PDB ID 8d33), which encodes the mitochondrial DNA polymerase subunit gamma-1, is an example of an AD/AR_mixed_ gene with a reported DN mechanism of pathogenesis. Missense variant positions are shown for the dominant (PEOA1) and recessive (PEOB1) forms of progressive external ophthalmoplegia, with their corresponding mLOF scores.

Another phenomenon that could explain the tendency for dominant and recessive variants in the same gene to be associated with similar phenotypes is the DN effect, as has been described in cases where DN variants phenocopy recessive disorders^30,35–38^. To test whether AD phenotypes with a predicted non-LOF mechanism have a tendency to phenocopy the recessive disorder, we analysed all non-redundant AD-AR phenotype pairs within AD/AR_mixed_ genes, representing 217 phenotype pairs in 103 genes. These were grouped into a high confidence non-LOF category if the mLOF score for the AD phenotype fell below the optimal threshold and was less than that for the AR phenotype. We then calculated the semantic similarity between AD-AR phenotype pairs using Human Phenotype Ontology^39^ terms, with the hypothesis that the non-LOF category should tend to have higher semantic similarity values due its enrichment in genuine DN mechanisms. As shown in **Figure 3B**, the non-LOF class displayed significantly higher AD-AR phenotype similarity values relative to other AD phenotypes (*P* = 9.2 × 10^-3^, Wilcoxon rank-sum test).

We further refined the analysis by filtering for genes whose DN-specific prior was greater than that for GOF and selecting the phenotype pair with the highest similarity for each gene. Pairs with a semantic similarity greater than 0.5 are listed in **Table 2**. Among these, *POLG* emerged as the top-ranking gene (**Figure 3C**). *POLG* encodes the mitochondrial DNA polymerase subunit gamma-1 and is associated with both dominant (PEOA1) and recessive (PEOB1) forms of progressive external ophthalmoplegia. These phenotypes share many clinical features, as implied by their disease names and semantic similarity score. Segregation studies have demonstrated that heterozygous LOF mutations in *POLG* are fully compensated by the normal allele^40^. Additionally, the dominant PEOA1-associated variant p.Tyr955His has been functionally characterised, demonstrating a DN effect^41^. These findings highlight the utility of combining mLOF scores with semantic similarity to identify DN disease phenotypes in mixed-inheritance genes.

**Table 2.**
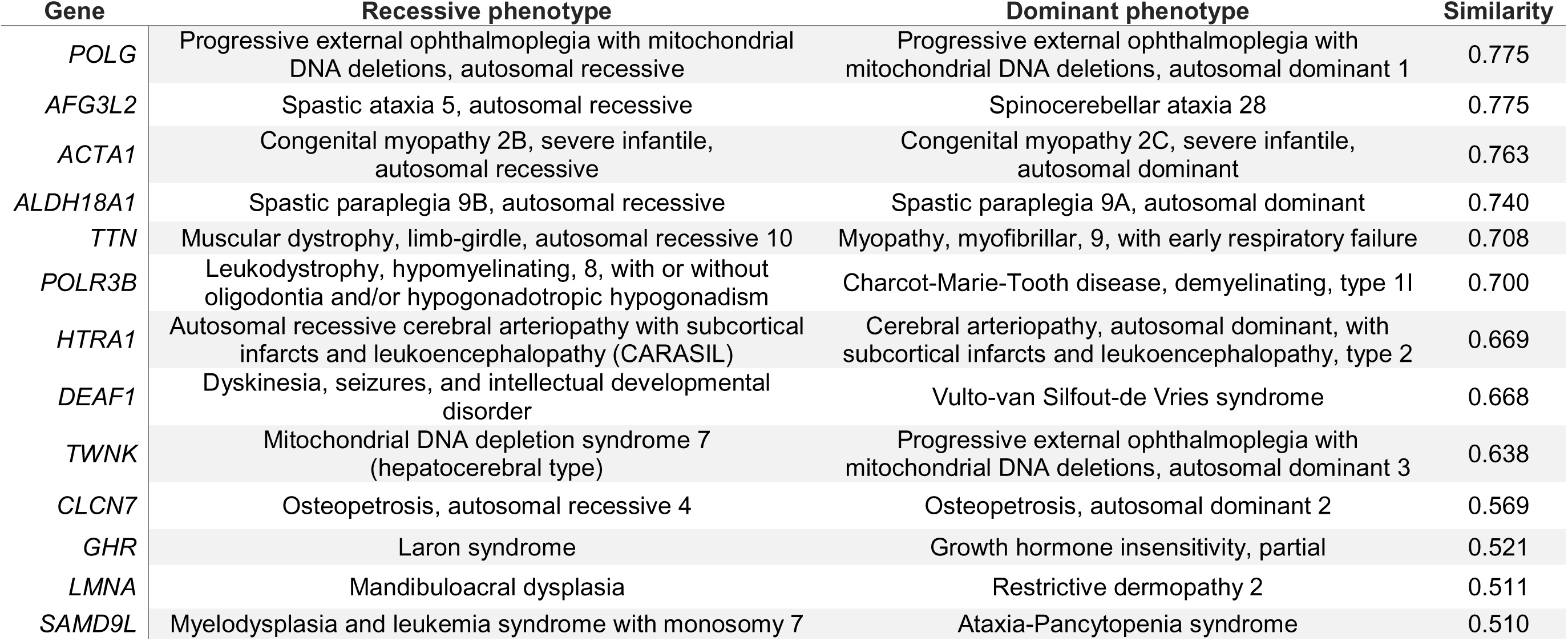
Top AR-AD phenotype pairs with high semantic similarity, where the dominant phenotype is likely to involve a dominant-negative effect.

### Disease phenotypes linked to distinct molecular mechanisms within the same gene

Given that mLOF score analysis suggested considerable mechanistic heterogeneity among multi-phenotype genes, we aimed to identify phenotype pairs most likely to exhibit distinct molecular mechanisms. To this end, we calculated the difference in mLOF scores for all possible phenotype pairs within multi-phenotype genes, excluding those with only recessive inheritance. We further refined our analysis by selecting pairs from beyond the 95^th^ percentile of the distribution, which we consider particularly interesting, and where one phenotype scored above and the other below the optimal threshold. **Table 3** summarises these genes, listing their phenotypes with higher mLOF scores (LOF-like) alongside those with lower mLOF scores (non-LOF-like). For some of the top-ranking genes, discussed in more detail below, protein structures and missense variant positions linked to the different phenotypes are shown in **Figure 4**.

**Figure 4.**
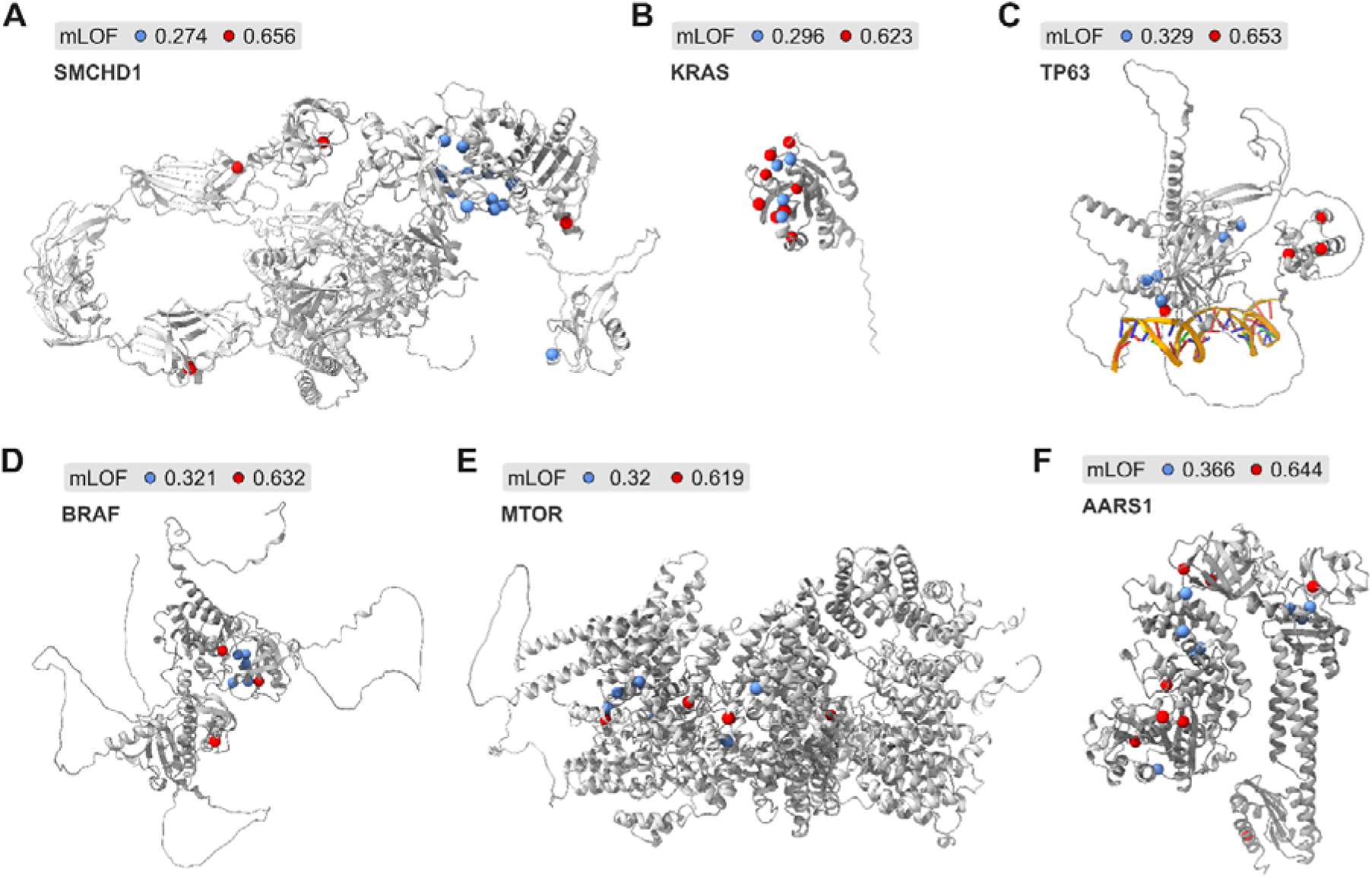
Examples of proteins having two disease phenotypes with mLOF scores indicating both loss-of-function and alternative mechanisms. Phenotype pairs in top ranking genes; see main text for a discussion on these and **Table 3** for their phenotype definitions. Structures are predicted models from the AlphaFold database. For TP63, the AlphaFold model was superimposed onto the DNA-bound TP63 complex (PDB ID 3us0). Red and blue spheres represent missense variants associated with the LOF-like and non-LOF-like phenotypes, respectively.

**Table 3.**
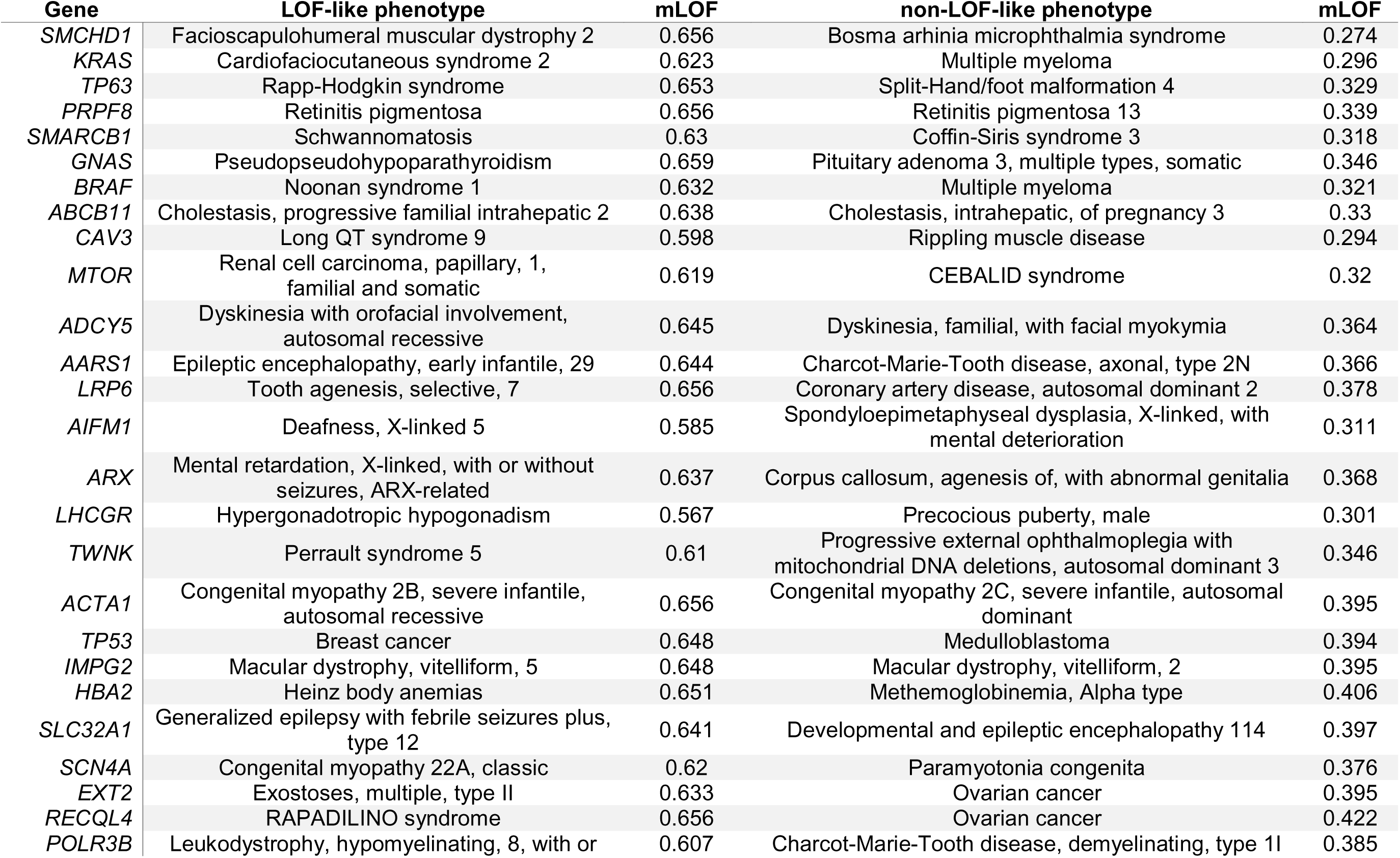
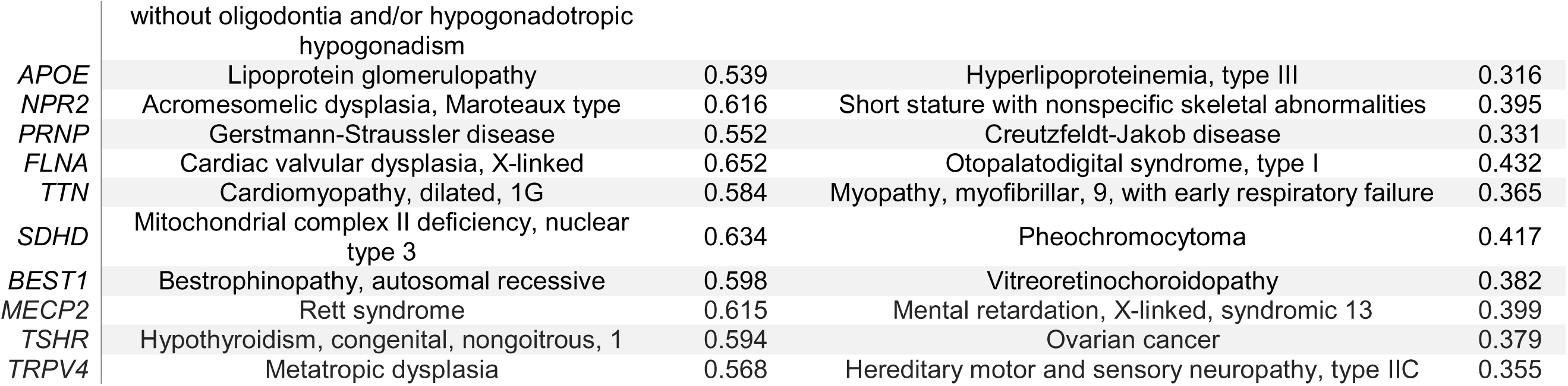
The most different phenotype pairs within multi-phenotype disease genes by the mLOF score.

SMCHD1 (**Figure 4A**) is a member of the structural maintenance of chromosomes protein family, which plays an essential role in epigenetic silencing. Mutations in the gene are linked to two distinct clinical phenotypes: the digenic dominant facioscapulohumeral muscular dystrophy type 2 (FSHD2) and the dominant Bosma arhinia microphthalmia syndrome (BAMS). In FSHD2, missense LOF mutations in SMCHD1, combined with a permissive D4Z4 haplotype on chromosome 4, lead to ectopic expression of DUX4, which is toxic to skeletal muscle cells^42^. Conversely, BAMS, characterised by the absence of the nose and accompanied by ocular and reproductive defects, is thought to result from GOF mutations^43^. Structural modelling revealed that BAMS-specific mutations cluster on the protein surface, pinpointing a cryptic interface^44^, a finding later confirmed by the crystal structure of the ATPase domain^45^. These observations are borne out by the high mLOF score of BAMS (0.656) and the low mLOF score of FSHD2 (0.274).

KRAS (**Figure 4B**) is a signalling protein and established oncogene with GTPase activity. The two phenotypes identified through mLOF analysis are cardiofaciocutaneous syndrome 2 (CFC2) and multiple myeloma. CFC2 is characterised by a distinctive facial appearance, heart defects, and intellectual disability^46^. Heterozygous missense variants underlying the phenotype are dispersed in the protein and have a highly structurally damaging effect, reflected by an mLOF score of 0.623. Supporting this, functional studies on the CFC2-associated variant p.Lys147Glu revealed weak GTP binding, falling short of the oncogenic threshold^47^. In contrast, multiple myeloma variants, which are typically highly recurrent somatic variants^48^, cluster around the GTP-binding site and are structurally mild, with an mLOF score of 0.296. Consistent with this, multiple myeloma is strongly linked to KRAS GOF variants^49^.

TP63 (**Figure 4C**) is a transcription factor required for limb formation from the apical ectodermal ridge^50^, linked to two dominant phenotypes: Rapp-Hodgkin syndrome (RHS) and split-hand/foot malformation 4 (SHFM4). RHS is characterised by anhidrotic ectodermal dysplasia and cleft lip and/or palate, and it is associated with LOF mutations in the sterile alpha motif domain (SAM)^51,52^. SHFM4, attributed to GOF mutations^51^, presents with clefts in the hands and feet, webbed fingers and toes, underdeveloped bones, and sometimes involving cognitive impairment. In agreement with their reported mechanisms, we found RHS to have a high mLOF score (0.653) due to strongly damaging mutations in the SAM domain, and SHFM4 to have a low mLOF score (0.329) as a result of much milder mutations at solvent-exposed residues.

Because TP63 forms tetramers via its oligomerisation domain^53^, and may form extended polymeric structures mediated by its SAM domain^54^, these structural features could suggest an assembly-mediated GOF (dominant-positive^1^) effect underlying SHFM4. For example, one SHFM4-associated mutation, p.Ala354Glu, is located in a region responsible for interacting with HIPK2^55^, which phosphorylates TP63 in response to DNA damage^56^.

BRAF (**Figure 4D**) is a serine/threonine-protein kinase and an established oncogene in human cancer^57^. Mutations in BRAF are linked to several clinical phenotypes, notably Noonan syndrome 1 (NS1) and multiple myeloma. Missense variants associated with NS1 have an mLOF score of 0.632, suggesting a LOF mechanism. These variants tend to be less clustered but more structurally damaging, and present with cardiac defects, facial dysmorphia, and reduced growth^58^. In contrast, missense mutations linked to multiple myeloma exhibit more activating effects, exemplified by the highly recurrent cancer-driver p.Val600Glu^58,59^. Multiple myeloma variants show a lower mLOF score of 0.321, likely reflective of an underlying GOF mechanism. These variants tend to be milder and localised exclusively within the kinase domain, a region critical for activating downstream signalling in the RAS-MAPK pathway.

MTOR (**Figure 4E**) is a serine/threonine protein kinase and the master regulator of cellular metabolism. mLOF score analysis has identified renal carcinoma and CEBALID syndrome (an acronym for craniofacial defects, dysmorphic ears, structural brain abnormalities, expressive language delay, and impaired intellectual development) to have missense variants with dissimilar effects on protein structure. Variants linked to renal carcinoma are dispersed across protein domains and are energetically impactful, yielding an mLOF score of 0.619. In contrast, CEBALID syndrome variants tend to be structurally milder and cluster near the ATP-binding site in the FATC domain, with an mLOF score of 0.32. GOF variants in MTOR have been previously linked to conditions such as Smith-Kingsmore syndrome^60^ and there is a growing body of evidence further implicating MTOR in developmental disorders^61–64^, with a recent *de novo* enrichment analysis detecting a significant missense burden in a cohort of in 31,058 parent-offspring trios^65^. Given that two MTOR subunits co-assemble into the mTORC1 complex, these mutations may exert DN or dominant-positive effects, potentially contributing to the observed phenotypic spectrum in MTOR-associated disorders.

AARS1 (**Figure 4F**) is the cytoplasmic alanine-tRNA ligase. mLOF score analysis revealed two distinct disease phenotypes: the recessive developmental and epileptic encephalopathy 29 (DEE29) and the dominant Charcot-Marie-Tooth disease, axonal, type 2N (CMT2N). Variants associated with DEE29 predominantly map to the ATP-binding site or the acceptor site recognition domain, consistent with its established biallelic LOF mechanism^66^. This is further supported by an mLOF score of 0.644, reflecting the severe structural impact of DEE29-associated mutations. By contrast, CMT2N variants are primarily located in the anticodon-binding domain and in a region homologous to the dimerisation interface observed in a remote paralogue^67^. These variants are associated with a lower mLOF score of 0.366, in agreement with their milder structural effects. Supporting this further, recent studies employing a humanised yeast assay suggest that missense variants linked to CMT2N exert a DN effect^68^.

### Mechanism prediction Google Colab notebook

To facilitate mLOF score calculation, we created a Google Colab notebook, available at https://github.com/badonyi/mechanism-prediction, allowing users to input a gene name or UniProt^69^ accession number along with a list of variants. The variants should map to the UniProt reference sequence—any mismatch between the variant and the reference amino acid sequence will be flagged with a warning. When only genomic variants are available, we recommend using the ProtVar^69^ web server to map these directly to the UniProt canonical isoform. We employ precomputed ΔΔG_rank_ values for the proteome and structures from the AlphaFold database^70^ to calculate EDC for the input variants. Although the latter limits proteins to <2700 amino acids, only about 1% of human proteins exceed this length. The results include all intermediary metrics, such as EDC and ΔΔG_rank_ values, the mLOF and the mechanism-specific posterior scores, as shown in **Figure 5**. A brief summary of the results is also provided to assist users in interpreting and reporting their findings.

**Figure 5.**
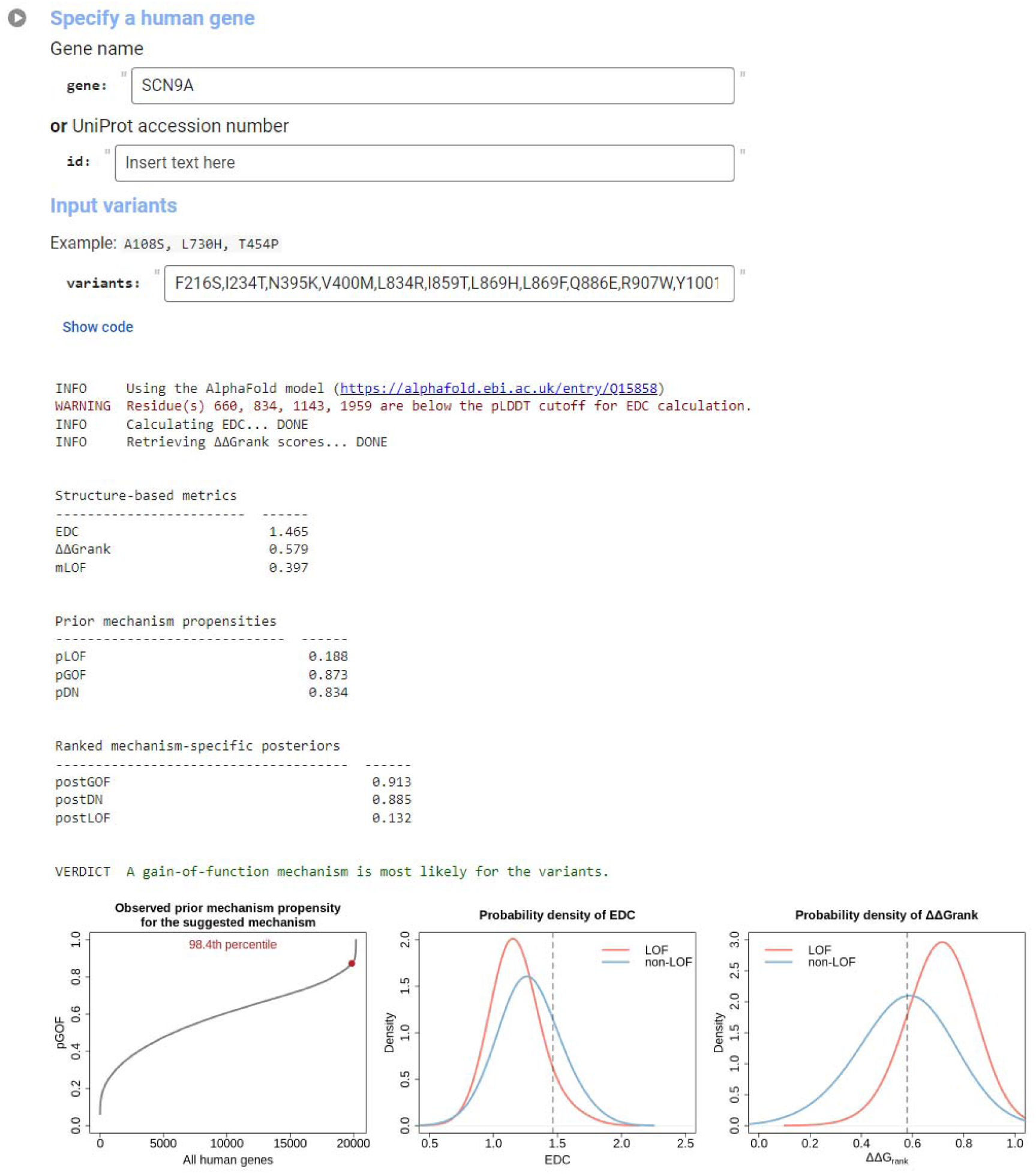
Example input/output of the Colab notebook. The notebook predicts molecular mechanisms for missense variants by accepting a gene name or UniProt ID along with a list of comma-separated variants in one-letter notation. Results include a link to the gene’s AlphaFold model used for EDC calculation, as well as the EDC, ΔΔG_rank_, mLOF score, and prior/posterior mechanism scores. The mechanism with the highest posterior score is suggested. Three plots are generated: (1) where the gene’s prior for the suggested mechanism lies relative to all human genes, and (2-3) where the observed EDC and ΔΔG_rank_ values fall within the empirical distributions of LOF and non-LOF genes.

## Discussion

Here, we developed an empirical distribution-based approach to calculate the missense LOF score, mLOF, which represents the likelihood that a group of pathogenic missense mutations will act via a simple LOF mechanism. We achieved this by leveraging fundamental protein structural properties of missense variants, their energetic impact (ΔΔG) and spatial clustering (EDC), both of which have an established and robust relationship to molecular disease mechanisms^15,18,21,24,71^. This approach offers two advantages. First, the use of a non-parametric kernel density estimation method preserves data interpretability at each step, allowing the use of intermediary results for hypothesis testing. Second, the applicability of ΔΔG and EDC to any combination of missense variants provides an optimal metric for assessing the missense LOF likelihood at the phenotype level. Variants within the same gene contributing to the same phenotype are more likely to be causally and functionally coupled, enhancing the precision of molecular mechanism predictions.

In our data, a quarter of genes whose disease phenotypes are linked to missense mutations have more than one associated phenotype. Although this proportion may be skewed by study bias, in that multi-phenotype genes are overrepresented in disease genes that have historically attracted more attention (*e.g.*, *TP53*, *KRAS*, and *BRCA2*), mLOF score analysis indicates that 43% of dominant and 49% of mixed-inheritance multi-phenotype genes exhibit phenotypes driven by both LOF and non-LOF mechanisms. This finding has important implications for the design of clinical trials and the development of therapeutic interventions, suggesting that in many cases, different disease phenotypes of the same gene may require distinct treatment strategies tailored to the underlying mechanism.

Many dominant phenotypes in mixed-inheritance (AD/AR_mixed_) genes are likely attributable to DN effects rather than simple LOF. While this is expected—given that the presence of recessive inheritance reduces the likelihood of haploinsufficiency^29^, and a GOF mechanism is unlikely to mimic a recessive disorder^1^—it is nonetheless valuable information from a clinical point of view. By combining mLOF scores with phenotype semantic similarity, we could prioritise phenotypes resembling the recessive disorder in the same gene, identifying cases that may result from DN mechanisms. However, this analysis was not feasible for genes where the same phenotype is inherited in both dominant and recessive patterns (AD/AR_same_). In such cases, challenges remain in determining which variants are pathogenic only in the homozygous state and whether dominant variants are as likely to exert DN effects as those in AD/AR_mixed_ genes.

In many mixed-inheritance genes, the distinction between dominant and recessive modes of action is clear: missense DN variants in *ITPR1* are associated with Gillespie syndrome^30^, whereas only recessive null variants have been linked to a clinically identical phenotype^72^. In other cases, however, the distinction is less straightforward. Both recessive and dominant missense mutations in *IGF1R* cause resistance to insulin-like growth factor 1, manifesting in intrauterine growth retardation; for example, the recessive p.Arg40Leu^73^ and the dominant p.Val629Glu^74^. Several genetic factors can explain this, *e.g.*, hypomorphic homozygous or compound heterozygous mutations (recessive partial LOF) that produce phenotypes indistinguishable from those caused by haploinsufficiency (dominant complete LOF). Another factor may be incomplete penetrance^75–78^, which causes variable phenotype expressivity within ancestries^76^, sexes^79^, and even families^80^. For example, the presence of common variants may modify the penetrance of inherited rare variants, lowering the liability threshold required for being affected by a disease^81^. Alternatively, DN variants might present as perfect phenocopies of the recessive disorder. In some possibly rare cases, a single variant could result in biallelic LOF, while exhibiting a DN effect in the heterozygous state, as suggested by emerging evidence in *AARS1*^82^.

Considering exclusively dominant genes, diverse genetic and mechanistic factors can explain the coexistence of haploinsufficient and DN phenotypes in the same^83^. For example, allele-specific expression and sex differences in *SMC1A* lead to distinct phenotypes, with truncating mutations linked to haploinsufficiency causing a seizure disorder and DN missense mutations resulting in Cornelia de Lange syndrome^84^. Notably, mutant SMC1A proteins maintain a residual function in males but confer a DN effect in females^85^. A similar phenomenon occurs in *NF1*, where truncating mutations reduce protein levels via haploinsufficiency, while destabilising missense mutations induce a DN effect by promoting degradation of the wild type in a tissue-specific pattern^86^. Although such contrasting mechanisms are scarcer among phenotypes induced only by missense mutations, these examples highlight the nuanced relationship between mutation type, biological context, and the resulting phenotype, complicating the interpretation of molecular mechanisms.

Unlike other molecular mechanism predictors, such as LoGoFunc^11^ or VPatho^12^, which are typically trained on mechanism labels using supervised learning approaches and may suffer from inflated performance estimates due to circularity issues^87^, the mLOF score is a simple, empirical metric independent of existing mechanism classifications. Moreover, the mLOF score also differs in functionality from these predictors. First, it relies on missense variants already known to be causal for a disease or have evidence supporting their involvement in pathogenesis, for example, through family studies or cohort sequencing. Second, rather than inferring the mechanism for a single variant, it harnesses collective properties of variants, which potentially makes its estimate more robust for variants related through their shared phenotype.

Despite the utility of mLOF scores, several limitations remain, many of which could be addressed as more structural data become available. First, our method is necessarily restricted to missense variants, because it relies on EDC and ΔΔG, measures that do not readily apply to other mutation consequences like stop-gain, indel, or frameshift variants with respect to differences between molecular mechanisms. Future efforts should focus on developing structure-based methods to evaluate these mutations in a mechanistic context, expanding our variant interpretation ability beyond missense variants. Second, for EDC calculation our method includes variants only within regions with a pLDDT (predicted local distance difference test) score^88^ above 70, limiting it to non-disordered and well-predicted regions of protein structure. Although pathogenic missense mutations are highly enriched in structured regions^89^, this limitation excludes certain variants, *e.g.*, those in short linear motifs^90^, from the scope of the mLOF score.

Our approach also assumes that the missense variants used to calculate the mLOF score are causal. This limits its direct application in patient cohorts where often multiple variants in the same gene are identified, and it is difficult to know which variants, if any, can be linked to the disease. In such cases, additional variant prioritisation methods, *e.g*., the use of variant effect predictors, are required before the mLOF score can be applied.

While structurally mild pathogenic variants often localise to functional sites, such as an interface, they are not exclusively non-LOF. For example, the mutation p.Ile87Arg in PAX6 leads to loss of DNA binding^91,92^, causing an aniridia phenotype through haploinsufficiency. Rarely, LOF variants may cluster, as seen in follicular lymphoma-associated mutations in EZH2, which concentrate in the SET domain, disrupting S-adenosyl-L-methionine binding^21^. Conversely, in few cases, DN variants may be highly structurally damaging. For instance, missense variants linked to late-onset retinal degeneration in C1QTNF5 occur in the C1q head domain, responsible for functional activity, while assembly into a trimeric complex is driven by a separate collagen-like domain. This physical separation may allow co-assembly of wild-type and functionally impaired subunits, despite structural damage in the C1q domain^93^.

Finally, while we currently rely on monomeric AlphaFold models for EDC and ΔΔG estimation, incorporating predicted structures of protein complexes^94–99^ could substantially improve the accuracy of missense LOF likelihood estimation by providing more biologically representative structural insights through the consideration of intermolecular interactions^15,100^. Furthermore, predicting subunits as part of a complex, rather than in isolation, often yields more accurate conformations due to the presence of buried contacts^101^, potentially making the spatial clustering of pathogenic residues more sensitive and informative.

In summary, we developed a broadly applicable and readily interpretable metric of missense LOF likelihood, the mLOF score. Our Google Colab notebook offers an accessible platform to compute the score and apply it within a Bayesian framework to predict to most likely mechanism for any combination of pathogenic missense variants in a human gene. This flexibility enables a deeper investigation into the structural effect of mutations, facilitating applications in variant interpretation and molecular mechanism studies.

## Methods

### ClinVar mapping to UniProt reference proteome

Genomic coordinates of ‘pathogenic’ and ‘likely pathogenic’ missense variants, which we refer to as pathogenic, were extracted from the ClinVar^22^ variant calling file (accessed 10-Sep-2024) using BCFtools^102^. These were subsequently mapped to human reference sequences in UniProt^69^ release 2024_04 with Ensembl VEP 112^103^. We retained phenotype cross-references to the OMIM database, as they represent the most comprehensive and reliable phenotype annotations. These were further annotated with MIM identifiers from two additional sources: the protein-specific JSON files with UniProt variation data via the EBI Proteins API^104^, and the UniProt index of human variants curated from literature reports (2024_04 of 24-Jul-2024). Gene-level inheritances were obtained from the OMIM database (06-Aug-2024) as previously described^17^. To obtain inheritance modes at the phenotype level, we accessed the phenotype_to_genes.txt and phenotype.hpoa files from the Human Phenotype Ontology (HPO) database^39^ (13-Aug-2024), which contain MIM identifiers and their HPO ontology terms. This process resulted in 63,920 pathogenic missense variants, of which 45,888 (71.8%) have an associated OMIM phenotype with an inheritance annotation, as summarised in **Figure S4**.

### Structural data

We computed EDC and ΔΔG_rank_ based on the predicted human structures downloaded from the AlphaFold database^70,105^ (AFDB). For the most part, we used AFDB v1 structures, which are consistent with the 2021_02 UniProt release. Any reference sequence between the lengths of 50 to 5,000 amino acids that has undergone a sequence change from the 2021_02 to the 2024_04 UniProt releases were remodelled with AlphaFold2, using the default settings of LocalColabFold (ColabFold^106^ v1.5.5) on an NVIDIA A100 GPU with 500 GB of RAM. Structures were visualised using UCSF Chimera X version 1.8^107^.

EDC was calculated as previously described^15^. For each residue, we determined the alpha carbon distance to pathogenic residue positions (‘disease’) and to all other (‘non-disease’) positions, keeping the shortest distance. The final metric is the ratio of the common logarithm of average non-disease and disease distances. Residues with pLDDT < 70 were removed from the calculation, because pathogenic missense mutations are highly enriched in structured regions^89^, therefore mutations in disordered proteins with a small structured core may appear clustered relative to the total volume of the model. For proteins modelled as multiple overlapping fragments in the AFDB, we took the fragment with the highest number of missense variants.

To compute ΔΔG_rank_, FoldX 5.0 was first used to estimate the change in the Gibbs free energy for all amino acid substitutions possible by single nucleotide change based on human codon usage. The RepairPDB command was called on each model before running the BuildModel command to estimate the ΔΔG. For pathogenic variants that map to multiple fragments of the same protein in the AFDB, we took the mean ΔΔG. The output values were ranked and rescaled, so that 0 represents the mildest mutation in the structure, and 1 the most damaging. Finally, for any group of variants (*e.g*., that belonging to a specific phenotype) we average ΔΔG_rank_ values to obtain the mean ΔΔG_rank_ metric, which we refer to as ΔΔG_rank_ for brevity. We note that raw FoldX ΔΔG values are available for non-disordered and well-predicted regions in human AlphaFold models via the ProtVar API^108^. However, as these do not allow calculation of ΔΔG_rank_, we have made our values available at https://osf.io/g98as.

### mLOF calculation

We use genes from Gerasimavicius et al.^15^ to fit our model, as these genes have at least one missense variant (rather than, *e.g.*, a protein truncating variant) associated with a molecular disease mechanism. At both the gene and phenotype levels, we require at least three missense variant positions with a pLDDT > 70 to ensure reliable estimates for EDC. We perform Gaussian kernel density estimation separately on the EDC and the ΔΔG_rank_ values of pathogenic missense variants in LOF and non-LOF genes, evaluating at 1024 equidistant points with three times the Sheather-Jones bandwidth^109^. The adjustment factor of three was chosen because, at this value, the probability distributions are smooth and monotonic without noticeable fluctuations, as shown in **Figure S1A-B**. To prevent extreme values from disproportionately influencing the probability estimates, we cap the empirical distributions at the 10^th^ and 90^th^ percentiles. We then compute the density functions for both groups and identify the closest point in the density function to each observation, allowing us to derive the estimated density values. *P*_LOF_(EDC) and *P*_LOF_(ΔΔG), which represent LOF probabilities of observed EDC or ΔΔG_rank_ values, respectively, are computed by dividing the density value of the LOF group by the sum of LOF and non-LOF density values. The combined estimate, *i.e*., the mLOF score, is obtained by taking the case-specific weighted mean of the two probabilities, which is considered a robust method when the dependence between the variables is strong or unknown^110^. *P*_LOF_(EDC) is weighted by the number of variants used for ΔΔG_rank_ calculation, while *P*_LOF_(ΔΔG) is weighted by the number of residue positions used for EDC calculation. This approach, which we refer to as the ‘weighted mean’ method, effectively weakens the influence of ΔΔG when variants are localised to disordered regions, thereby strengthening that of EDC. The previous steps are visually represented in **Figure S1**. Finally, to estimate a posterior mechanism likelihood score, we use pDN, pGOF, and pLOF from our proteome-scale model as informed priors, which reflect the likelihood of observing the given mechanism when missense variants are identified in a gene^18^. These priors are updated with the mLOF score according to Bayes’ rule. We formalise our probabilistic framework below:

*Definitions:*

Let *x* be a single observation of EDC or mean ΔΔG_rank_.

Let *cap_LOF_* and *cap_non-LOF_* be the cap values for the observations.

Let *x*’ be the capped observation.

Let *f_LOF_*(*x*’) and *f_non-LOF_*(*x*’) be the density functions at observation *x*’.

Let *d*_*points*_ be the vector of points where the density functions are evaluated.

Let *index* be the index of the closest value in the density function.

Let *w*_ΔΔG_ be the number of unique residue positions used for EDC calculation.

Let *w_EDC_* be the number of variants used for mean ΔΔG_rank_ calculation.

*Cap values for a given observation X*:

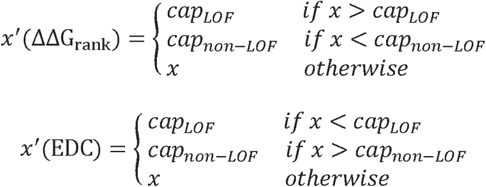

*Finding indices of nearest density points for a given capped observation X* ^I^:

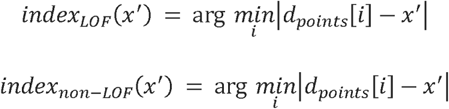

*Obtaining density values*:

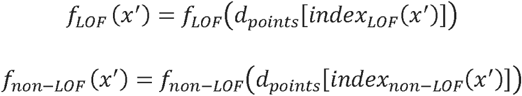

*Calculating P_LOF_(EDC)* and *P_LOF_*(*ΔΔG*):

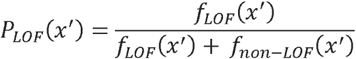

*Calculating the mLOF score:*

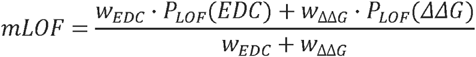

*Calculating mechanism-specific posterior scores:*

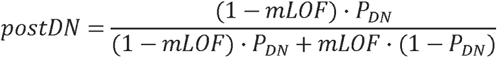

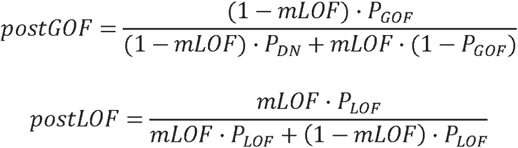

### Method validation

We initially compared the performance of the weighted mean method to a generalised linear model (GLM) that estimates mLOF from EDC and ΔΔG_rank_ and an interaction term between them. The rationale was that a GLM may better capture the joint distribution of the metrics, potentially outperforming the weighted mean method, which relies on marginal distributions. By comparing bootstrapped AUROC estimates, we found that the posterior mechanism-specific scores obtained with the GLM-based mLOF score had a consistently worse performance across the binary class pairs. A possible explanation for this is that mLOF scores from the GLM (**Figure S1D**) are much less conservative than those from the weighted mean model (**Figure S1C**), leading to the mLOF score having a greater influence on the posterior. This result suggested that a generalised linear model cannot achieve the same performance as our weighted mean model, supporting its use.

To evaluate the mLOF score’s utility in distinguishing different molecular mechanisms within the same gene, we applied it to DN, GOF, and LOF variants identified by high-throughput functional assays in TP53^27^ and HRAS^28^. For TP53, we adopted the classification approach described in the original study: DN and LOF variants were defined as those with Z-scores three standard deviations (SD) away from the mean of all synonymous mutations, based on the ‘p53^WT^+nutlin-3 assay’ for DN variants, and on the ‘p53^NULL^+nutlin-3’ and ‘p53^NULL^+etoposide’ assays for LOF variants. For visualisation in **Figure S3A**, a combined scored was calculated as (‘p53^WT^+nutlin-3’ + ‘p53^NULL^+nutlin-3’ – ‘p53^NULL^+etoposide’) / 3. For HRAS, we defined LOF variants as those with relative enrichment values falling more than two SD below the mean in the ‘DMS_regulated’ assay and GOF variants as those exceeding two SD above the mean in the ‘DMS_attenuated’ assay. We note that this classification is more stringent than the one SD approach used by the authors. For the purpose of visualisation in **Figure S3B**, the combined score was calculated as (‘DMS_regulated’ + ‘DMS_attenuated’ + ‘DMS_unregulated’) / 3.

For the analysis with LoGoFunc-predicted GOF and LOF variants^11^, we downloaded genome-wide missense variant predictions from the GOF/LOF database (release 10-Aug-2024, https://itanlab.shinyapps.io/goflof/). Genomic coordinates of these variants were mapped using the Ensembl VEP 112 pipeline (as described above) and merged with our ClinVar dataset. We computed the mean LoGoFunc_GOF score for variants associated with phenotypes in single-phenotype AD genes and evaluated it against all genes in our GOF *vs* LOF dataset, as well as the corresponding test set. The test set excludes genes used for training the model (used to construct pGOF) and is limited to proteins with <50% pairwise sequence identity.

### Phenotype-level analyses

To ensure reliable mLOF estimates, we considered disease phenotypes with missense variants at 3 distinct positions. In **Figure 2A-B**, the following criteria was used to create the phenotype classifications. Note that gene-level molecular mechanisms are based on our previous study^18^.

1. **AR**: the gene is exclusively AR in OMIM and the phenotype is annotated as AR in HPO.
2. **[AR]-AD/AR_mixed_**: the gene has at least one AD and one AR phenotype with a sufficient number of missense variants, and the phenotype is annotated as AR in HPO.
3. **[AD]-AD/AR_mixed_**: the gene has at least one AD and one AR phenotype with a sufficient number of missense variants, and the phenotype is annotated as AD in HPO.
4. **AD/AR_same_**: the phenotype has both AD and AR inheritance annotation in HPO.
5. **XLR**: any phenotype of genes in OMIM with ‘X-linked recessive’ or ‘X-linked’ inheritance.
6. **AD**: the gene is exclusively AD in OMIM and the phenotype is annotated as AD in HPO.
7. **LOF**: the phenotype is annotated as AD in HPO, with the gene either having a reported LOF mechanism or has ‘Sufficient evidence for dosage pathogenicity’ in the ClinGen database^34^ as of 10-Sep-2024. Does not overlap with DN or GOF genes.
8. **GOF**: the phenotype is annotated as AD in HPO and the gene has a reported GOF disease mechanism. Excludes AD/AR_mixed_ genes. May overlap with DN genes.
9. **DN**: the phenotype is annotated as AD in HPO and the gene has a reported GOF disease mechanism. Excludes AD/AR_mixed_ genes. May overlap with GOF genes.
10. **Unknown**: the gene is exclusively AD in OMIM, lacks a reported disease mechanism, and the phenotype is annotated as AD in HPO.

For each within-gene phenotype pair, we calculated how missense variant sets relate to each other in terms of overlap: (1) distinct, if the variant sets are mutually exclusive; (2) intersect, where some variants are shared between the sets; (3) subset, if variants of one phenotype represent a subset of the other; and (4) identical, if the variant sets are mutually inclusive. **Figure S4C-D** illustrate the relative proportions of set relationships across all non-redundant phenotype pairs and within unique inheritance groupings. As the mLOF score can be affected by the extent of variant overlap, losing discriminatory value for ‘identical’ sets, we only considered phenotype pairs whose variants had ‘distinct’ and ‘intersect’ set relationships.

Semantic similarity between AD-AR phenotype pairs was calculated with the ontologyIndex and ontologySimilarity R packages based on the 08-Feb-2024 HPO release, using Lin’s expression of term similarity^111^.

### Statistical analysis

Data analysis was performed in R 4.3.0^112^, using the tidyverse metapackage. Statistical tests were two sided and an alpha level of 0.05 was considered significant. In bootstrap analyses, 1,000 resamples were used. The optimal threshold was derived by selecting the value that minimises the combined Euclidean distance from the (0,1) coordinate of the ROC curve, based on the true positive and false positive rates. In box plots, boxes represent data within the 25^th^ and 75^th^ percentiles, the middle line is the median, the notches contain the 95% confidence interval of the median, and the whiskers extend to 1.5× the interquartile range.

## Data and code availability

Data and analysis code are available at https://osf.io/ah2uc. **Table S1** can be downloaded at https://osf.io/29pxr. Precalculated ΔΔG_rank_ values for all missense variants in the human proteome can be downloaded at https://osf.io/g98as. The Colab notebook for mechanism prediction can be accessed here: https://github.com/badonyi/mechanism-prediction.

## Acknowledgements

This project was supported by funding from the European Research Council (ERC) under the European Union’s Horizon 2020 research and innovation programme (grant agreement No. 101001169) and by funding from the Medical Research Council (MRC) Human Genetics Unit core grant (MC_UU_00035/9). JAM is a Lister Institute Research Prize Fellow. This work has made use of the resources provided by the Edinburgh Compute and Data Facility (ECDF).

---

**Figure S1.**
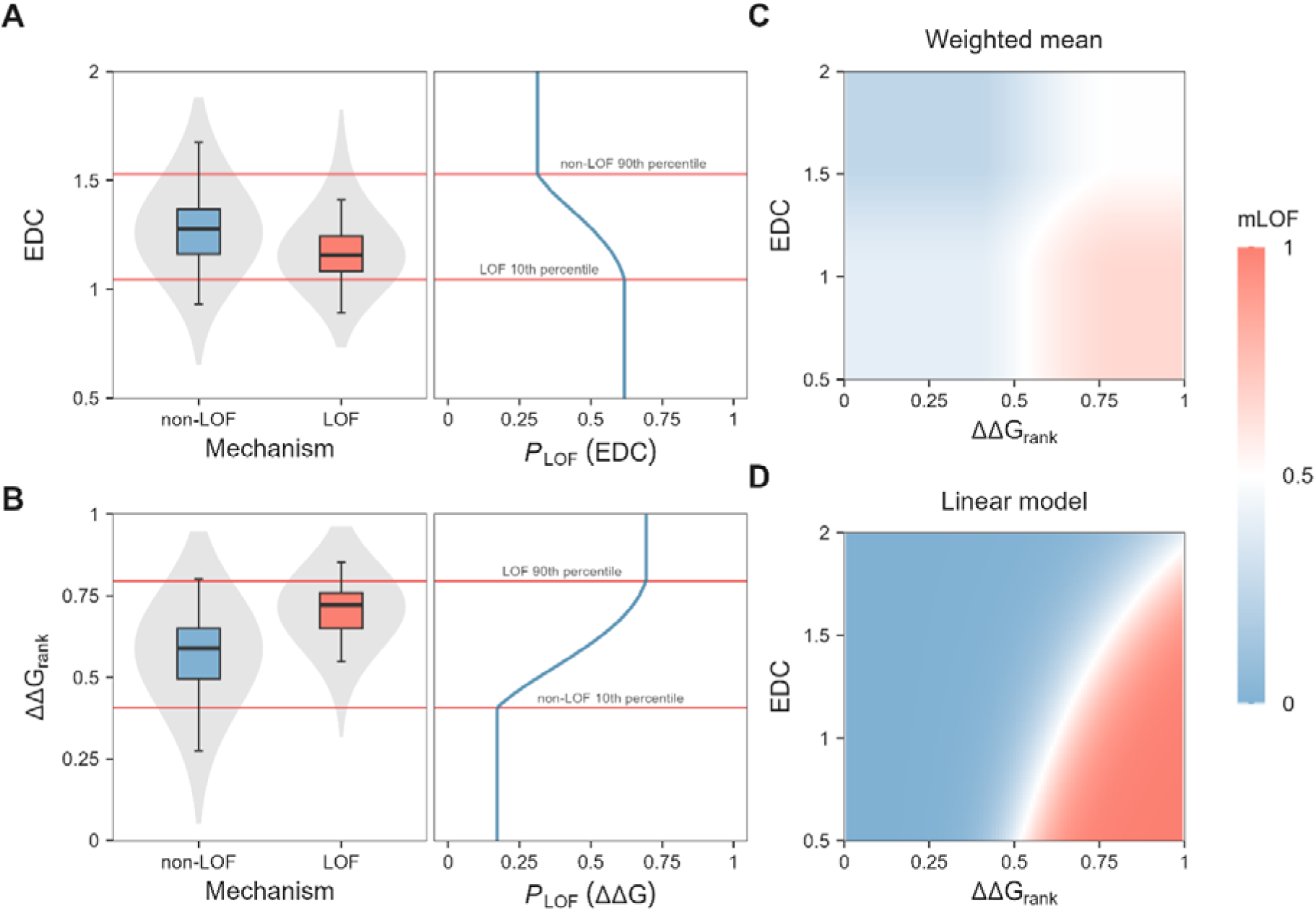
An empirical distribution-based approach to derive the mLOF score. **A**: EDC and **B**: ΔΔG_rank_ distributions of genes primarily associated with missense LOF and non-LOF mechanisms. The plot on the right shows the distribution of estimated LOF probability given the observed EDC or ΔΔG_rank_ value (*P*_LOF_(EDC) and *P*_LOF_(ΔΔG), respectively). Red lines represent 10^th^ and 90^th^ percentile caps. **C**: Landscape of combined probabilities (mLOF score) for the expected range of EDC and ΔΔG_rank_ values with the weighted mean method (used in this study), using a global median weight of 1.47:1 (EDC:ΔΔG_rank_) for the purpose of this visualisation. **D**: Landscape of mLOF scores with the generalised linear model (not used in this study).

**Figure S2.**
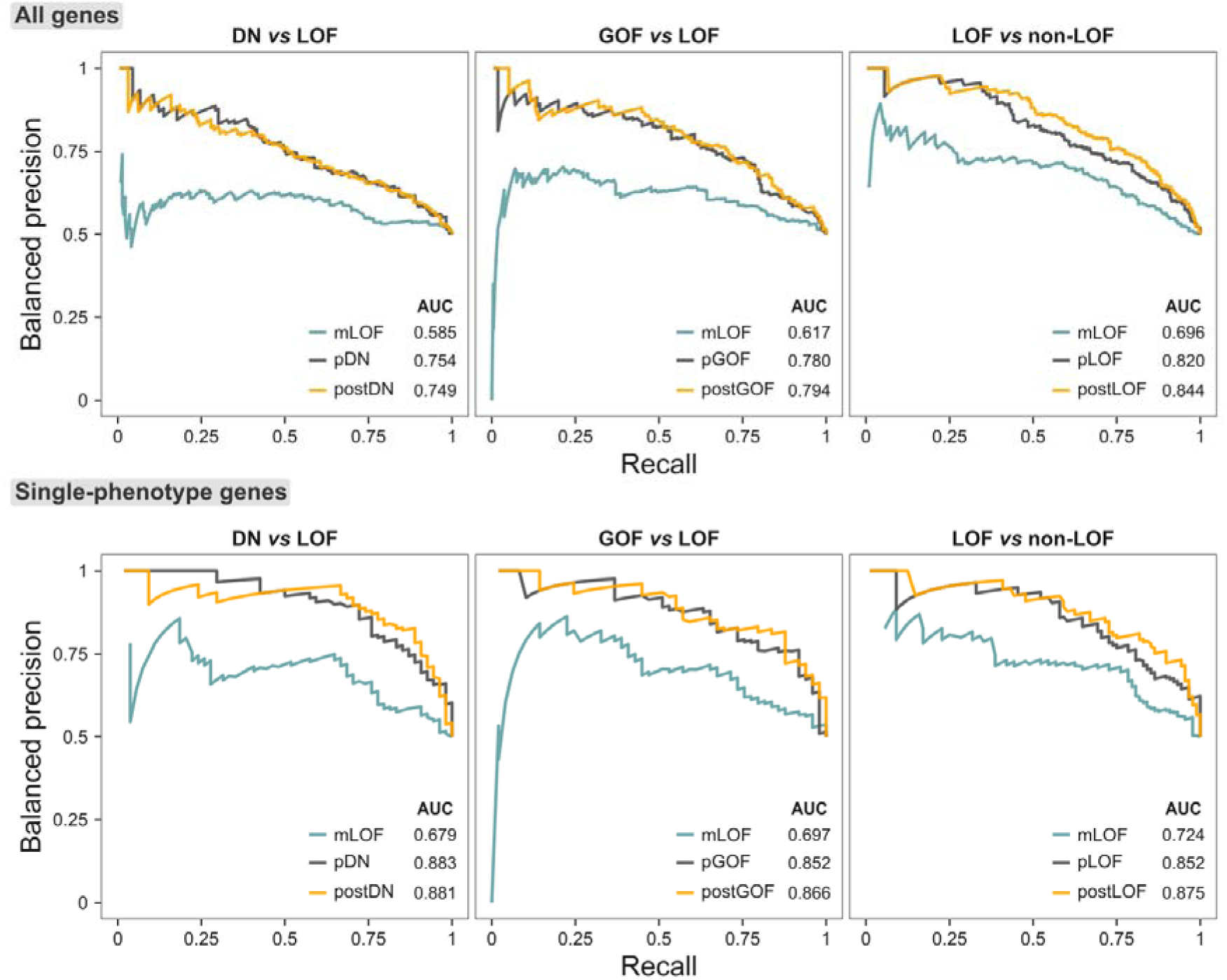
Balanced precision-recall curves related to. **Figure 1B**. Phenotype-level balanced precision-recall curves and area under the curve (AUC) values of the mLOF score, the prior mechanism probability for the gene (one of pDN/GOF/LOF), and the posterior mechanism-specific scores (one of postDN/GOF/LOF) across the binary class pairs used to construct the priors, split into all genes and single-phenotype genes.

**Figure S3.**
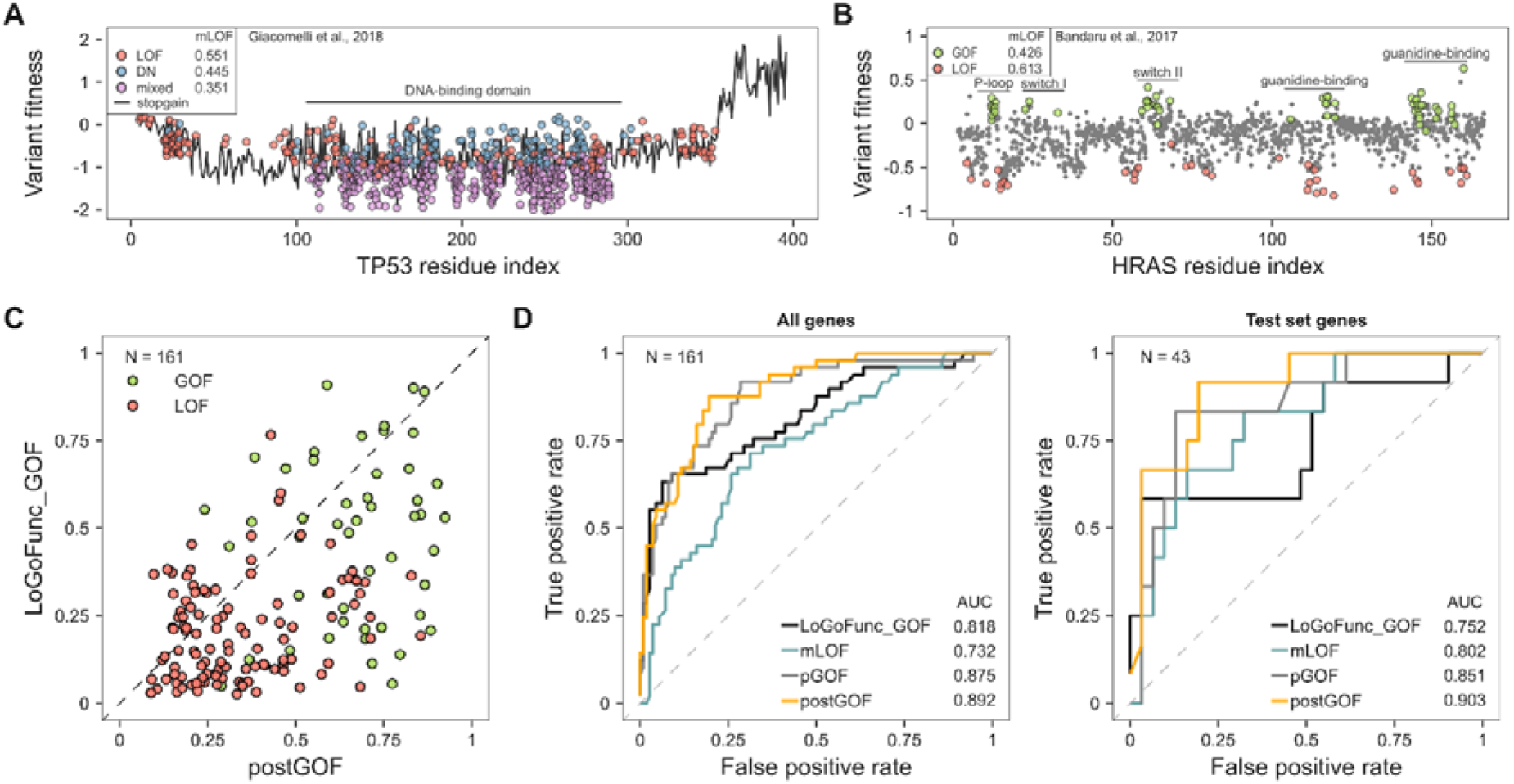
Comparison of the mLOF score with functional assay results and computational predictions of GOF variants. **A**: Missense LOF variants in TP53 detected by high-throughput functional assays^27^ have a higher mLOF score than missense DN variants. **B**: mLOF scores support the distinction between missense LOF and GOF variants in HRAS based on deep mutational scanning data^28^. **C**: Scatter plot of postGOF *vs* average LoGoFunc_GOF probability for phenotypes in single-phenotype AD genes, coloured by their reported GOF/LOF disease mechanisms. N is the number of phenotypes/genes. **D**: Receiver operating characteristic (ROC) curves and area under the curve (AUC) values of the mLOF score, pGOF, postGOF, and the average LoGoFunc_GOF probability for phenotypes. N is the number of phenotypes/genes.

**Figure S4.**
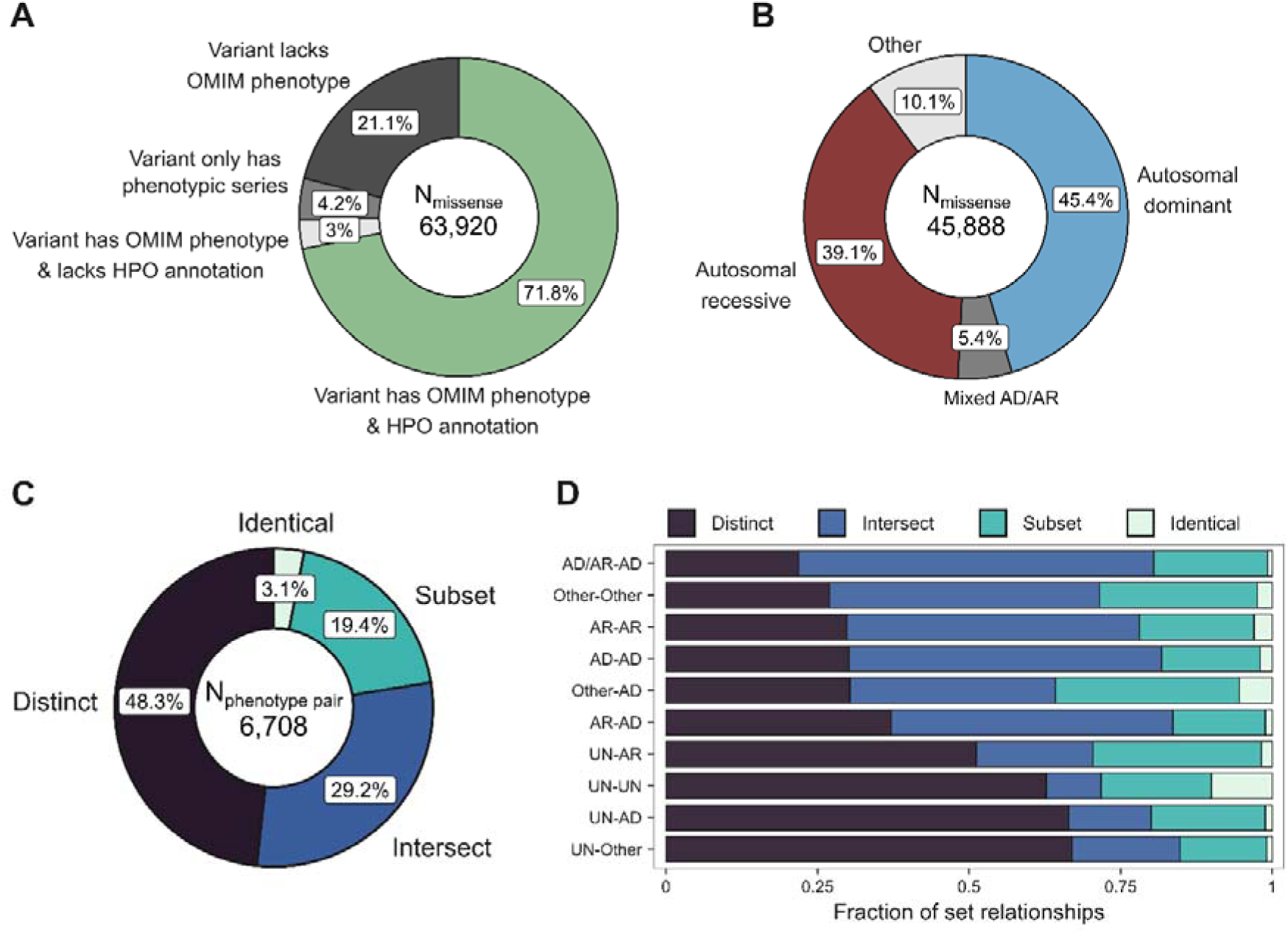
Missense variant statistics in terms of phenotype composition and overlap. **A**: The composition of available OMIM phenotype annotations for pathogenic missense variants after cross-referencing with HPO data. **B**: Inheritance composition of variants with OMIM phenotype & HPO annotation. **C**: Profile of within-gene inheritance pairs considering their shared missense variants. Non-redundant inheritance pairs with at least 100 occurrences are shown.

